# Capturing the spatiotemporal dynamics of task-initiated thoughts with combined EEG and fMRI

**DOI:** 10.1101/346346

**Authors:** Lucie Bréchet, Denis Brunet, Gwénaël Birot, Rolf Gruetter, Christoph M. Michel, João Jorge

## Abstract

When at rest, our mind wanders from thought to thought in distinct mental states. Despite the marked importance of ongoing mental processes, it is challenging to capture and relate these states to specific cognitive contents. In this work, we employed ultra-high field functional magnetic resonance imaging (fMRI) and high-density electroencephalography (EEG) to study the ongoing thoughts of participants instructed to retrieve self-relevant past episodes for periods of 20s. These task-initiated, participant-driven activity patterns were compared to a distinct condition where participants performed serial mental arithmetic operations, thereby shifting from self-related to self-unrelated thoughts. BOLD activity mapping revealed selective activity changes in temporal, parietal and occipital areas (“posterior hot zone”), evincing their role in integrating the re-experienced past events into conscious representations during memory retrieval. Functional connectivity analysis showed that these regions were organized in two major subparts of the default mode network, previously associated to “scene-reconstruction” and “self-experience” subsystems. EEG microstate analysis allowed studying these participant-driven thoughts in the millisecond range by determining the temporal dynamics of brief periods of stable scalp potential fields. This analysis revealed selective modulation of occurrence and duration of specific microstates in both conditions. EEG source analysis revealed similar spatial distributions between the sources of these microstates and the regions identified with fMRI. These findings support growing evidence that specific fMRI networks can be captured with EEG as repeatedly occurring, integrated brief periods of synchronized neuronal activity, lasting only fractions of seconds.

**Significance:** We investigated the spatiotemporal dynamics of large-scale brain networks related to specific conscious thoughts. We demonstrate here that instructing participants to direct their thoughts to either episodic autobiographic memory or to mental arithmetic modulates distinct networks both in terms of highly spatially-specific BOLD signal oscillations as well as fast sub-second dynamics of EEG microstates. The combined findings from the two modalities evince a clear link between hemodynamic and electrophysiological signatures of spontaneous brain activity by the occurrence of thoughts that last for fractions of seconds, repeatedly appearing over time as integrated coherent activities of specific large-scale networks.

## Introduction

The occurrence of spontaneous conscious thoughts is difficult to grasp experimentally (1). Spontaneous mentation is neither random nor meaningless (2), however how to precisely capture the wandering mind and attribute it to specific cognitive thoughts, is yet unclear. Functional magnetic resonance imaging (fMRI) as well as electroencephalography (EEG) studies have previously related resting-state activity to cognitive processes mainly indirectly by using post-scan questionnaires (3) or by asking participants about their ongoing thoughts during the spontaneous mentation (2, 4). The striking similarities found by comparing the spatial distributions of large-scale networks specific to self-reported thoughts and those initiated by experimental tasks (5, 6), contribute to the wide debate about the content of spontaneous mental activity at rest and the relation of resting state networks (RSN) to cognitive networks (7–9).

Time-resolved fMRI studies previously demonstrated that the RSNs spontaneously fluctuate in and out of spatially and temporally overlapping correlations (10–14), indicating that temporal dynamics of resting state activity are not stationary, but rather partitioned into distinct epochs. This observation corroborates the prevailing concept that spontaneous mental activity is discontinuous and parsed into a series of conscious states that manifest discrete spatiotemporal patterns of neuronal activity (for review see (15–17)). However, despite much progress in dynamic resting-state functional neuroimaging, the delayed and slow hemodynamic response to neuronal activity limits fMRI methods to capture these states (11). Indeed, to efficiently execute mental processes, large-scale networks have to dynamically re-organize on *sub-second* temporal scales (14, 18). EEG microstate analysis allows to investigate these fast temporal dynamics of large-scale neural networks and to access information about the functional organization of spontaneous mentation in time (17, 19). EEG microstates reflect brief epochs of synchronized activity during spontaneous mentation that persist for around 100 milliseconds (17, 20). Previous studies demonstrated that the occurrence and duration of EEG microstates determine the quality of spontaneous mentation, and as such could represent the basic building blocks of conscious mental processes (4). EEG microstates thus qualify as the electrophysiological segmentation of ongoing mental activity into short-lasting brain states (17, 21, 22). Yet, despite several efforts using simultaneous EEG-fMRI (23–25), the functional and neurophysiological relation between EEG microstates and fMRI resting states has not yet been fully resolved (17).

Recent studies have explored novel experimental designs based on task-initiated spontaneous activity, whereby participants were asked to think freely, but with specific instructions on the thoughts they should focus on (5, 6). These approaches have yielded promising insights into the cognitive underpinnings of spontaneous thoughts, and are well suited to the recording and analysis of continuous activity with techniques like EEG and fMRI. The overall goal of this study was to capture the occurrence of spontaneous thoughts within specific, externally-controlled cognitive domains, combining ultra-high field 7T fMRI and high-density 64-channel EEG recordings to obtain both hemodynamic and electrophysiological signatures with high functional sensitivity. Fifteen healthy participants were recorded during uninstructed spontaneous mentation (hereafter termed “rest condition”) and while focusing their thoughts repeatedly for periods of 20sec with eyes closed on either episodic, self-related memories associated with a briefly presented image (“memory condition”), or arithmetic calculations (“math condition”) (*SI Appendix,* Figs. S1). Consistent with cognitive tasks that require working memory and direct the attention outside of the self (26), we chose the math condition in order to selectively de-activate regions of the default mode network (DMN), which we expected to become active in episodic memory retrieval (5, 27).

To probe functional changes specific to the distinct cognitive domains, we compared the spatial distribution of local BOLD activity changes, as well as their organization in large-scale networks, across conditions. We then examined the fast temporal organization of the brain’s large-scale network dynamics, using the EEG microstate approach. Specifically, we focused on the most basic characteristics of the EEG microstate temporal dynamics: their duration, their frequency of occurrence and their transition probabilities. Finally, we estimated the brain networks generating the EEG microstates and compared them to the fMRI RSNs.

## Results

### Instructed thoughts modulate fMRI networks

To identify changes in spontaneous activity across different brain regions during the two stimulus-initiated thought conditions i.e. math and memory conditions, the fractional amplitude of low-frequency fluctuations (fALFF) (29) was mapped, serving as a model-free measure of local BOLD activity (30). Having obtained a fALFF estimate for each brain voxel of each fMRI run and participant, these values were then compared between conditions on the group level using topological false-discovery rate (FDR) inference (31) (T>2.20, α=5%). Condition-specific fALFF changes were thereby found for several cortical and subcortical brain regions, as follows:

The regions with significantly higher brain activity during the memory compared to the math condition included the core brain areas of the episodic memory retrieval network (32–34) in the parietal, the medial temporal, the prefrontal and occipital lobes (Fig. 1a). We found dominant activity in the lateral part of the parietal lobe: bilaterally the supramarginal gyrus (lrSMG, BA40) and the right angular gyrus (rAG, BA39) of the inferior parietal lobe (IPL). Activity was also found in the medial part of the parietal lobe: the left precuneus (lPCu, BA7), the dorsal part and the ventral part of the posterior cingulate cortex/retrosplenial cortex (dPCC, BA31; vPCC/RSc, BA23. In the temporal lobe we found bilaterally-increased activity in the parahippocampal gyri (lrPHG, BA28,35,36) and the right inferior temporal gyrus (rITG, BA21). Significant activity was also found in the right inferior frontal gyrus (rIFG, BA45), and bilaterally in the lateral occipital gyri (lrLOCG, BA19). When comparing the fALFF estimates in these brain regions to the non-instructed rest condition (considered as baseline) we found that some of the memory-math differences were due to a stronger increase in BOLD activity during the memory compared to math condition (areas included lrPHG and dPCC), while other changes were due to increased BOLD activity during the memory and decreased BOLD activity during the math conditions. Areas de-activated during math compared to rest included rAG, lrSMG, lrLOCG, lPCu and vPCC/RSc. (Fig. 1b). Conversely, we also identified a set of brain areas more active during the math compared to the memory condition that have been previously associated to mental arithmetic tasks and frontoparietal control network (FPCN) in general (35) (*SI Appendix,* Figs. S2a and S2b).

**Fig. 1.**
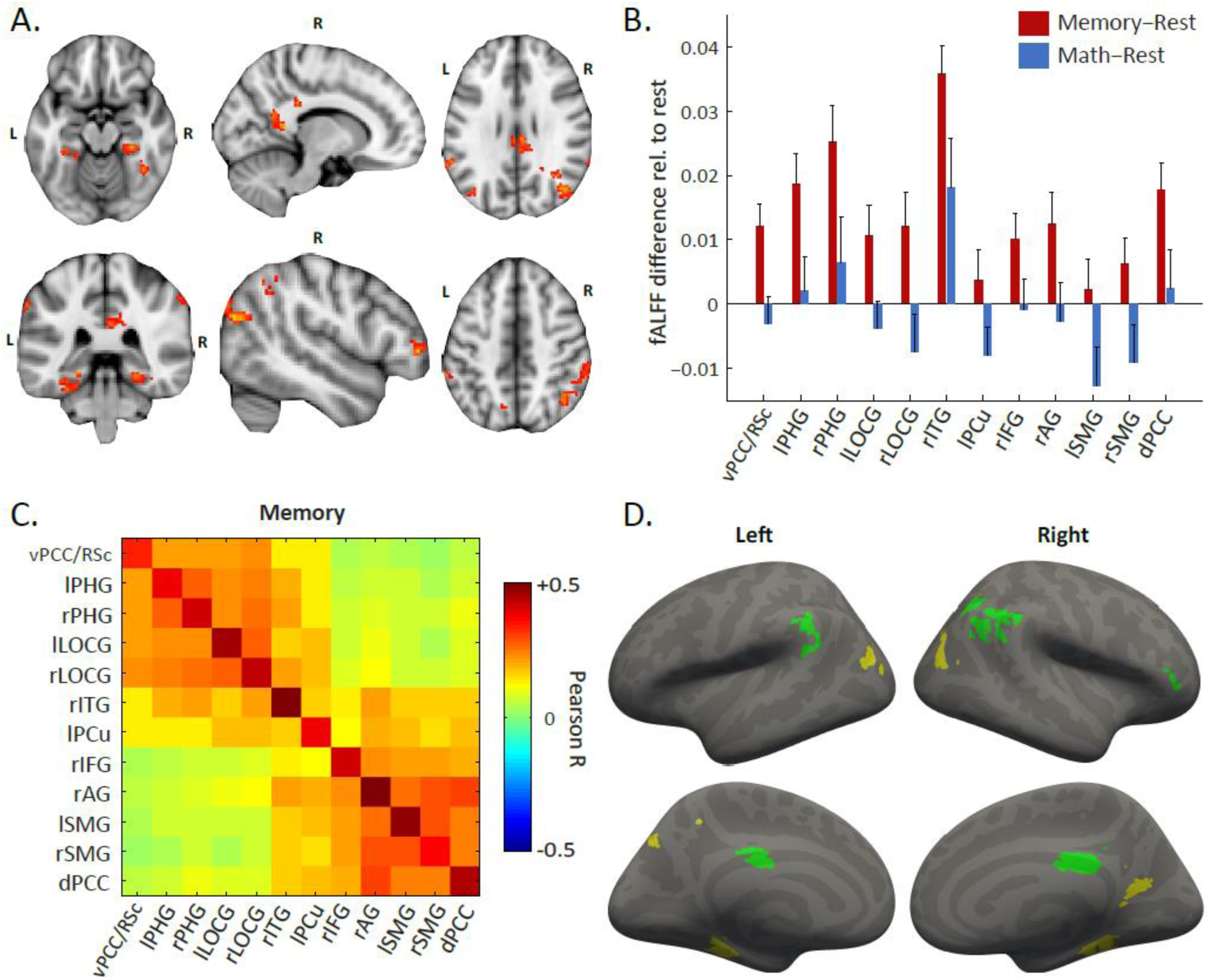
Memory-specific changes in fMRI activity and fMRI functional connectivity memory sub-networks. **A. fMRI fALFF analysis.** T-test of fALFF for memory > math condition revealed stronger activity in lrSMG, rAG, lrLOCG, lPCu, dPCC, vPCC/RSc, lrPHG, rITG, rIFG. T-score threshold: T>2.20; FDR-corrected for multiple comparisons (5%). **B. Areas of stronger activity in the memory, compared to math condition.** Some of the memory-math differences were due to a stronger increase in BOLD activity during the memory than the math condition relative to rest, while other changes were due to increased BOLD activity during the memory and decreased BOLD activity during the math conditions relative to rest. **C. Connectivity analysis.** Network I. included the vPCC/RSc, lrPHG, lrOCG, rITG and lPCu, which we called “scene-reconstruction subsystem”. Network II. comprised the rIFG, rAG, lrSMG, and dPCC, called “self-experience subsystem”. **D. Sub-networks of the connectivity analysis.** Regions were also projected onto a surface template for better visualization.

To determine condition-specific networks, we further analysed, in terms of their functional connectivity, the brain regions that demonstrated significant BOLD activity changes between the memory and math conditions, as identified by the fALFF analysis. These connectivity values were organized in matrices, and the regions were re-ordered according to their correlation profiles. The connectivity analysis revealed two distinct networks within the regions that had displayed stronger activity during the memory condition (Figs. 1c and 1d). Network I. was strikingly similar to a previously described subsystem of the DMN (5, 36), which becomes involved during constructions of mental scenes (hereafter termed “scene-reconstruction subsystem”). Network II. closely matched a previously defined subsystem of the DMN (5, 36), which becomes active when participants engage in self-relevant cognitive processes and reflect on their current mental states (hereafter termed “self-experience subsystem”). Compared to the non-instructed rest condition, both networks displayed strong increases in connectivity amidst their respective regions, while remaining uncorrelated to each other and to the areas identified in the math condition. Our results support the multi-component aspects of DMN, showing that different regions of this network behave differently and form functionally distinct subsystems based on the cognitive function involved (36, 37).

### Instructed thoughts modulated the temporal dynamics of specific EEG microstates

Having found a high spatial organization of functionally connected networks during the memory vs. math conditions using fMRI, we then examined the fast temporal organization of these large-scale networks, by identifying the most dominant EEG microstate maps for each of the three conditions (i.e. math, memory and rest conditions) using a *k*-means cluster analysis. The optimal number of clusters, determined by a meta-criterion (*SI Appendix,* SMethod 3) appeared to be six for each condition. The topographies of these six maps were strikingly similar between the three conditions and resembled those previously described in the literature (17). We ordered the microstate maps according to their highest spatial correlation across conditions (i.e. math, memory, rest) and labelled them from A-F states (17, 38) (Fig. 2a). Statistical comparison of the underlying brain sources that generated these microstates indicated similar networks between the corresponding maps in the three conditions (*SI Appendix,* Table S1). We then fitted the maps back to the original EEG of each participant and labelled each time point with the microstate that had highest spatial correlation (Fig. 2b). This procedure allowed us to determine the mean duration of each microstate in the three conditions, and how often the microstates occurred independent of their duration.

**Fig. 2.**
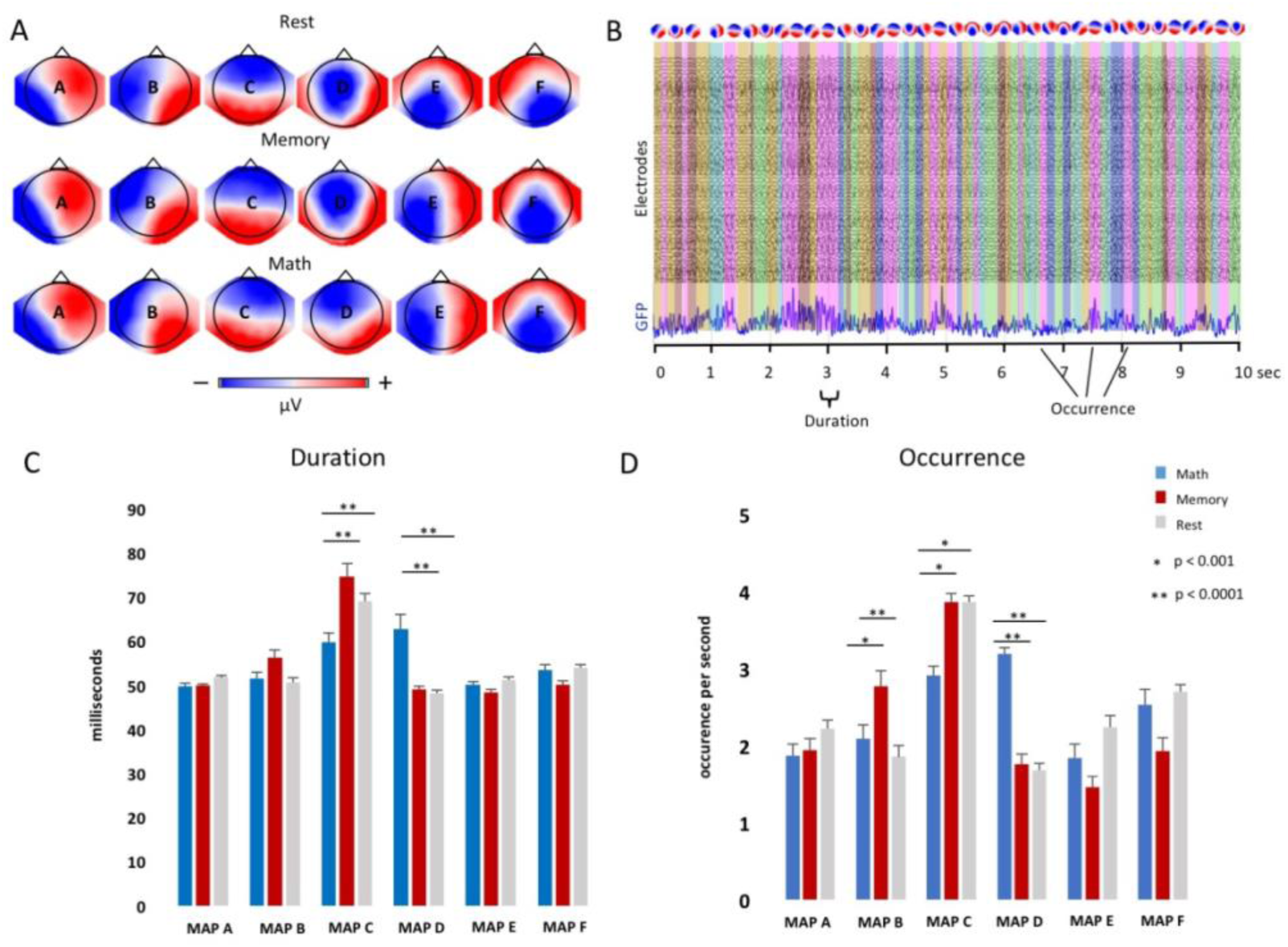
Temporal dynamics of EEG microstates. **A.** The six microstates identified by k-means cluster analysis across subjects in the three conditions (rest, memory and math). **B.** A representative period of 10-sec EEG with eyes closed after picture presentation is shown (traces of 64 electrodes and Global Field Power (GFP) trace). Back-fitting the 6 microstate maps derived from the cluster analysis shows the chunking of the EEG into segments of various durations covered by one of the microstate maps (indicated by different colors). **C.** Mean and standard error of the duration of each of the microstates in each condition. Post-hoc tests were performed when the ANOVA revealed significant condition differences at p<0.0001. This was the case for microstate map C and map D: Microstate C was significantly shorter and microstate D significantly longer in the math condition compared to memory and rest. **D.** Occurrence of the six microstates in the three conditions (number of microstates per second). Significant ANOVAs were found for microstates B, C, and D. As for the duration, microstate C occurred less and microstate D more often in the math condition compared to rest and memory. Concerning microstate B, an increased occurrence was found for memory compared to math and rest.

ANOVA analyses for each microstate revealed that two states (i.e. microstates C and D) significantly differed in their duration and occurrence depending on the condition (Fig. 2c) and one state (i.e. microstate B) significantly differed in its occurrence (Fig. 2d). Microstate B was specific to the memory condition as it significantly increased in occurrence compared to the math as well as to the rest conditions (both p<0.0001). Microstate C was significantly decreased in duration and occurrence in the math condition as compared to both, the memory and the rest conditions (both p<0.001). Finally, microstate D was specific to the math condition as it significantly increased in occurrence and duration compared to the memory (p<0.001) and the rest (p<0.0001) conditions.

### Sources of EEG microstates correspond to the fMRI RSN

To estimate networks underlying each microstate we inverted the original data of each subject into source space using a distributed linear inverse solution (39), as described in the Methods. We then averaged the estimated activity across all time points that were labelled with the same microstate maps for each subject and each condition. The brain regions underlying the six microstate maps confirmed and extended previous efforts (38) in source localization of microstates (Fig. 3). Specifically, microstate A showed left-lateralized activity in the superior temporal gyrus (STG), the medial prefrontal cortex (MPFC) and the OCG. Microstate B showed main activity in OCG and in the medial part of the parietal cortex in the PCu/RSc. The sources of microstate C were located bilaterally in the lateral part of the parietal cortex including both the SMG and AG. The sources of microstate D showed main activity bilaterally in the IFG, dACC, and superior parietal lobule (SPL)/intraparietal sulcus (IPS). Strongest activity for microstate E was found in the right MPFC. Finally, microstate F showed bilateral activity in the MPFC.

**Fig. 3.**
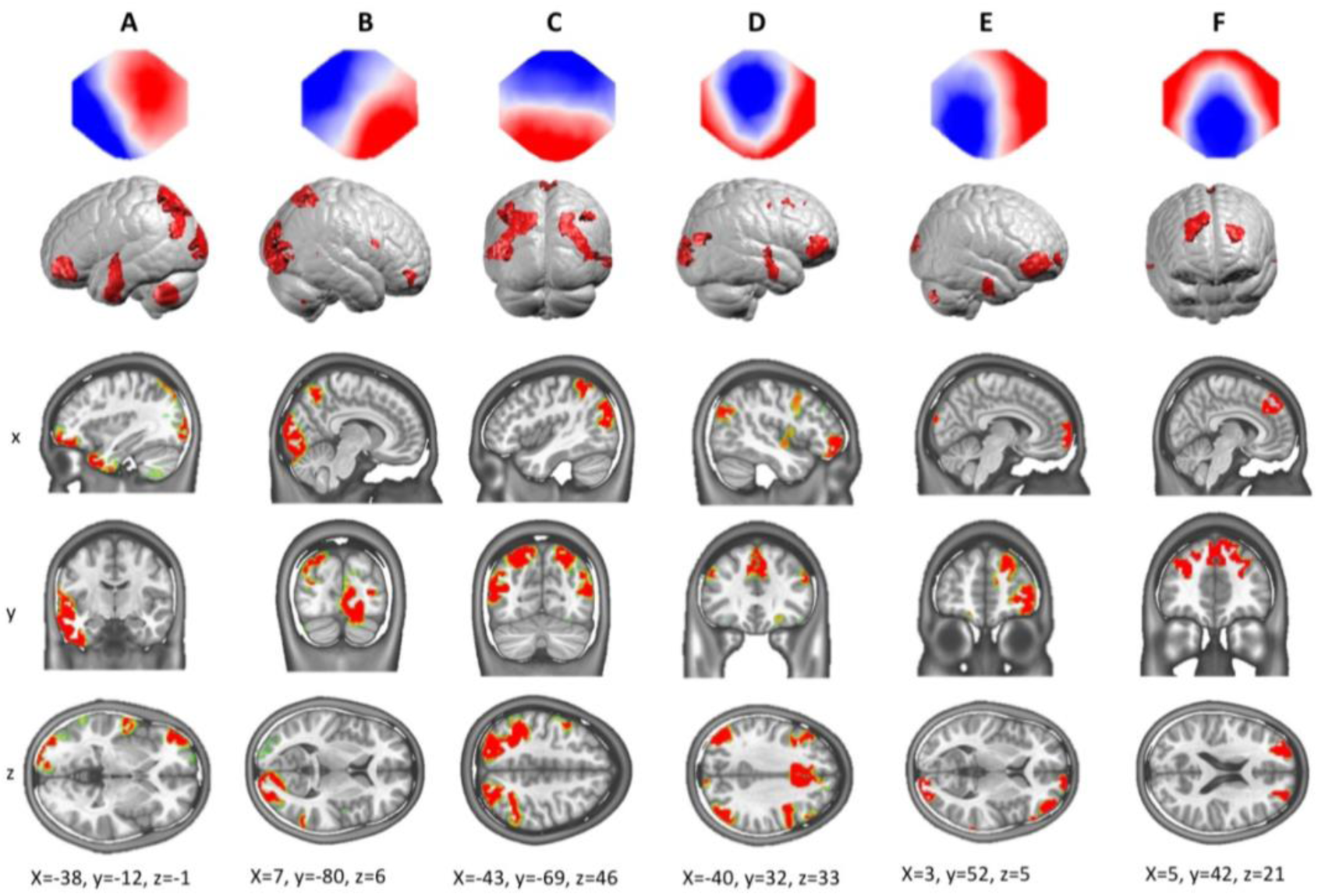
Source localization of EEG microstates. The EEG of each participant and condition were subjected to a distributed linear inverse solution and standardized across time. The source maps of all time points that were labelled with the same microstate map were then averaged within participants. The mean sources across subjects in the memory condition are illustrated here (for individual source maps per condition see *SI Appendix,* SFigs. 3-5). Areas with activity above 95 percentiles are shown. Notice the strong activity of the superior parietal lobe for microstate C (strong presence in memory and rest) and the frontoparietal activity for microstate D (strong presence in math).

Remarkably, the network underlying microstate C, which occurred more often and lasted longer in the memory compared to the math condition, largely overlapped with the lateral parietal areas of the “self-experience” memory retrieval subnetwork that we identified in the fMRI analyses. On the other hand, the brain areas underlying microstate B, which selectively increased in occurrence during the memory condition, overlapped with the medial parietal areas of the “scene-reconstruction” subnetwork that we detected in the fMRI analyses. Finally, brain areas underlying microstate D, which selectively increased in duration and occurrence in the math condition, included areas generally attributed to the frontoparietal control network (FPCN) (40).

### Increased transition between memory-related subnetworks

To determine whether the brain alternates between the two dominating microstates in the memory condition reflecting networks that are involved in mental scene reconstruction (microstate B) and self-relevant cognitive processes (microstate C), we calculated the transition probabilities (normalized by the occurrence) between microstate C and D and compared them to all other possible transitions. We found that specifically and only in the memory condition, the transition from microstate C to microstate B was significantly more frequent than the transition from any other state and that microstate B transitioned significantly more often to microstate C than to any other state (Fig. 4). Likewise, microstate C was significantly more often followed by microstate B than microstates A, D or E, but not F.

**Fig. 4.**
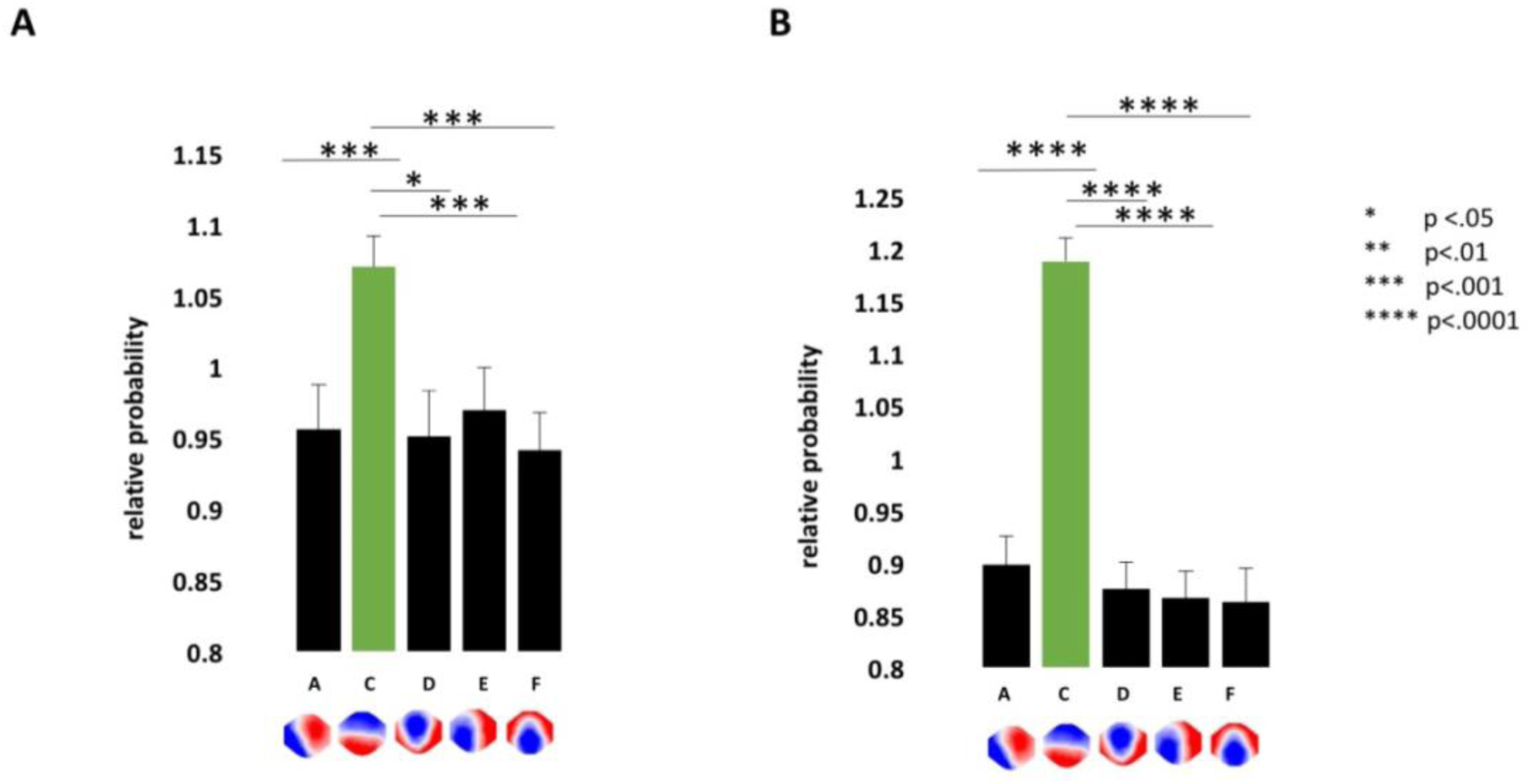
Markov chain transition probabilities. We calculated the transition probabilities from each microstate to any other using Markov chains. The observed probabilities were divided by the expected probabilites to account for the variability in occurrence of the states. In the memory condition, we found that microstate B was significantly more often preceded **(A)** and followed **(B)** by microstate C than any of the other microstates. This transition behavior was not found during rest or when participants performed mental arithmetic operations.

## Discussion

By combining fMRI and EEG in a paradigm where participants were instructed to focus their thoughts on specific tasks, we here provide direct evidence of capturing specific large-scale brain networks that are involved in self-related and self-unrelated thoughts. Particularly, the fMRI connectivity results revealed two memory subsystems, the “self-experience” and “scene-reconstruction” networks with distinct functional and anatomical characteristics. While this result is in line with previous task-related fMRI studies (36, 41) and confirms the effectiveness of this type of “instructed mentation” paradigm to reveal thought-specific networks, the EEG microstate analysis revealed new insights in the temporal dynamics of these networks in the sub-second range. We found that specific EEG microstates, representing brief periods of synchronized network activity, were selectively increased in duration and occurrence by the instructed thoughts. The brain regions generating these microstates overlapped with the specific fMRI networks. Moreover, during memory retrieval, we found increased transitions between the microstates that corresponded to the “self-experience” and “scene-reconstruction” subnetworks in the fMRI. These results provide direct evidence that the RSNs captured by fMRI are tightly linked to, and possibly originated by, a prolongation and repeated occurrence of states of synchronized activity of specific large-scale neuronal networks, in the sub-second timescale.

To capture the occurrence of any conscious experience, and to directly investigate the cognitive processes operating during mind-wandering, it is crucial to better control the spontaneous thoughts of participants (42, 43). In line with other recent studies (5, 6, 44), we here initiated periods of spontaneous mentation with brief presentations of external stimuli and instructed participants to close their eyes and internally direct their thoughts to either self-related photographs or self-unrelated arithmetic operations. The main regions that showed increased BOLD activity in the memory condition comprise the IPL and MTL structures. Numerous studies have previously identified these regions as the core of the episodic memory retrieval network (32–34). Traditionally, since the discovery of densely amnesic patients, the MTL structures have been regarded as essential for long-term memory formation, allowing us to remember past experiences and to retrieve acquired knowledge (45, 46). In contrast to the MTL structures, extensive evidence from patient lesion and brain stimulation studies suggests that the IPL plays a key role in integrating vivid details of personally experienced events into conscious representations during memory retrieval (47, 48). The subjective experience of perceiving a scene, recognizing a face, hearing a sound or reflecting on the experience itself presents a complex interplay between memory, attention and consciousness (49, 50). Tulving (51) defined the underlying ability to re-experience the subjective sense of self in the past and to mentally project oneself into the future, i.e. the autonoetic consciousness, as the crucial aspect of episodic memory retrieval. Evidence across lesion studies, stimulation and recording studies consistently point to the posterior regions, including temporal, parietal and occipital areas (“posterior hot zone”) as playing a direct role in specifying the contents of consciousness (43). Our current findings are also in line with a recent study (52) where we confirmed the contribution of the IPL, especially the AG, to the subjective, first-person perspective re-experience of self-relevant, vivid past episodes.

Our next aim was to capture the sub-second changes in temporal dynamics of brain states during the memory or math-initiated spontaneous mentation. Brain activity constantly fluctuates in and out of different mental states that are stable for fractions of seconds. Only one epoch or state of conscious content can be considered at a time (53). It is assumed that the EEG microstates capture these states that last for around 100 milliseconds only (21, 22). Using high-density whole-brain EEG, we indeed observed modulations of duration, occurrence, and transitions between particular microstates by the instructed thoughts, microstate B and C being increased in the memory condition and microstate D being increased in the math condition.

The topographies of these three microstates strongly resemble three of the four canonical microstates previously described in the literature (for reviews see (17, 20)). Studies on large cohorts showed that microstate C is generally the most dominant state during eyes-closed rest (54). Recently, Seitzman et al. (55) showed that microstate C decreases in duration and occurrence during a mental arithmetic task, similar to our findings. Decrease of microstate C duration has also been described when subjects are engaged in object or verbal visualization tasks compared to rest (56). Based on these and other studies, it is assumed that microstate C reflects activity in the DMN (38, 55-57); for a discussion, see (17). Indeed, our analysis of the sources underlying this microstate confirms such interpretation. The sources of microstate C were located bilaterally in the lateral part of the parietal lobe and MTG, areas that we attributed in the fMRI results to the “self-experience subsystem” of the DMN. The observation that microstate C is not significantly boosted by the memory condition compared to rest is not surprising. It confirms the assumption that self-relevant memory retrieval also predominates during spontaneous mind wandering, hence the sub-system of the DMN including the lateral areas of the parietal lobe is strongly active.

Moreover, our investigation revealed that microstate B increased in occurrence during the memory as compared to both rest and math conditions. This state has been previously attributed to the visual network (23, 38). For example, Milz et al. (56) showed that microstate B increased when participants were asked to visualize previously presented images. Source localization in our study also indicated strong activation of the visual cortex together with medial parietal areas. Intriguingly, lesions to the medial parietal cortex cause memory recognition and visuospatial impairments, but no impairments related to self-consciousness (58). We thus interpret the increased occurrence of microstate B as related to the “scene-reconstruction subsystem” found in the fMRI connectivity analysis. An intriguing assumption is therefore that the microstate analysis allows us to disentangle the sub-parts of thoughts related to the conscious experience of an episodic autobiographic memory, i.e. visualization of the scene and visualization of the self in the scene. Indeed, the transition probability analysis revealed a more frequent switching between microstate B and C than between any other state.

Additionally, our results show that while both microstates B and C are less frequently appearing when participants are engaged in math calculations, microstate D strongly increases in duration and occurrence during this condition. Microstate D has previously been attributed to the attention/cognitive control network including frontoparietal areas (23, 38) and our source localization of microstate D confirmed the attribution of this state to the frontoparietal control network (FPCN) (40). Furthermore, the observation of increase of microstate D and decrease of microstate C fits very well to the observation that the FPCN and the DMN are inversely activated when participants are engaged in external-directed and self-directed cognition (59). Therefore, we show that the large-scale network anti-correlation found in the fMRI data is associated with sub-second modulation of the presence of microstates sub-serving these functions.

EEG microstate studies have repeatedly revealed changes of the temporal dynamics of microstates in mental disease, particularly schizophrenia (for reviews see (17, 20)). The most robust finding, confirmed in a recent meta-analysis (60), is an increase in duration and occurrence of microstate C and a decrease of microstate D in patients with schizophrenia (61, 62) or at risk to develop schizophrenia (63, 64), a disequilibrium that is normalized when patients are treated with antipsychotic medication (65) and with rTMS (66). These findings correspond well to the interpretation that microstate C reflects introspective self-focused thoughts while microstate D reflects attention and cognitive control. An increase of microstate C and decrease of microstate D in schizophrenia might index the progressive detachment of mental states from environmental input. While a healthy person constantly and effortlessly balances periods of rest with periods of focused attention when interacting with their surroundings, patients with schizophrenia or other mental disorders may persist on thinking about a particular unpleasant event that involves themselves and lose control over the natural flow of the wandering mind.

Understanding the functional significance of microstates with studies as the one presented here might thus not only be relevant for monitoring the vulnerability of patients at risk for mental disease and the effects of treatment, but also for better understanding the thoughts that these patients are caught in.

## Materials and Methods

### Participants and Experimental Paradigm

We included 15 healthy participants (30.5±5.5years, 5 male/10 female) in this study. The work was approved by the institutional review board of the local ethics committee, and all participants provided written informed consent prior to the experiment.

The paradigm consisted of three distinct conditions during eyes closed: rest (6min), mentally retrieving personal past episodes (10min), and mental arithmetic (10min). Before the recordings, all participants performed a classical autobiographical memory questionnaire (ABMQ)(28) in order to select vividly remembered images for the memory retrieval condition. Before each of the memory retrieval and mental arithmetic trials, participants saw a personal image/calculation for 5sec, after which they closed their eyes and had to either retrieve the event or continually subtract the numbers (20sec). For more details see (*SI Appendix,* SMethod1).

### fMRI Acquisition and Analysis

The fMRI sessions were conducted on a 7T head scanner (Siemens, Germany), equipped with an 8-channel head RF array (Rapid Biomedical, Germany). For details see (*SI Appendix,* SMethod2). During each paradigm, functional data were acquired with TR = 1.0s and 2.0mm isotropic spatial resolution, simultaneously providing fairly high temporal and spatial resolution, with whole-brain coverage. An additional 5-volume scan was also performed with reversed phase encoding direction, for subsequent correction of EPI distortions. Whole-brain T_1_-weighted anatomical images were acquired to aid spatial co-registration, with 1mm isotropic spatial resolution.

The data were processed using the FMRIB Software Library (FSL v5.0, UK) combined with Matlab routines developed in-house (Mathworks, USA). The pre-processing included motion correction, slice-timing adjustment, brain segmentation, spatial smoothing (FWHM = 4mm), temporal de-trending (100s cut-off), and regression of motion-derived confounds. Each fMRI dataset was co-registered to the standard MNI space after B_0_-unwarping, and using the subject’s anatomical space as intermediate step for the linear registration.

Voxel-wise fractions of low-amplitude fluctuations (fALFF) (29) were estimated for the pre-processed fMRI data of each subject and paradigm, in native space, and then brought to MNI space for group analysis. These fALFF maps were compared between the memory and math conditions using paired T-tests across subjects, and statistical significance was determined using topological false-discovery rate (FDR) inference to correct for multiple comparisons (31) (T > 2.20, FDR α = 5%).

The brain regions (clusters) displaying significant fALFF changes between the memory and math conditions were further considered for connectivity analysis. These regions were warped to each individual run’s native space, and the functional connectivity between each pair of regions was estimated as the average Pearson correlation between the time course of every voxel in the first region with that of every voxel in the second region. The estimates were organized in a connectivity matrix, and also included an estimate for each region paired with itself, serving as a measure of functional homogeneity within the region.

### EEG Acquisition and Analysis

EEG was recorded with a 64-channel Brain Amp EEG system (Brain Products, Munich, Germany). Offline, the EEG was down-sampled to 250Hz, bad electrodes were interpolated using a 3-D spherical spline, band-pass filtered between 1-40Hz and re-computed to the common average-reference. Infomax-based Independent Component Analysis (ICA) was applied to remove oculomotor and cardiac artefacts (for details see *SI Appendix,* SMethod3).

The free academic software Cartool was used for the microstate analysis (67). It uses a modified k-means cluster analysis to determine the most dominant scalp potential maps (ignoring polarity) at time points of maximal Global Field Power (17). The optimal number of cluster maps was determined first individually and then on the group level using a meta-criterion (*SI Appendix,* SMethod 3). This cluster analysis resulted in 6 cluster maps for each condition.

The six group cluster maps of a given condition were fitted back to the original EEG by calculating the spatial correlation with the cluster maps and labelling each data point with the cluster map that showed the highest correlation (23). Two temporal parameters were then quantified for each recording of each subject: the mean duration of the continued presence of a given map, and the number of times per second a given microstate appeared – termed “occurrence” (Figs. 2c and 2d). To investigate differences between conditions and microstates we performed repeated measure ANOVAs with the factors “condition” (rest, math, memory) for each microstate class. We applied a threshold of p<.0001 to correct for multiple testing and performed post-hoc comparisons only if this threshold was reached.

Finally, we analysed the syntax of EEG microstates by computing the probabilities from a single *n*^*th*^-previous state to the current one for each subject and transition pair and normalized it by all between-class transitions (62).

In order to estimate the sources contributing to each of the six microstates we calculated a distributed linear inverse solution (LAURA)(39). The lead field for the inverse solution was calculated for 64 electrode positions and the average brain of the Montreal Neurological Institute in a grey matter constrained head model using the LSMAC head model with 5000 distributed solution points (67). A standardization across time was applied for each solution point in order to eliminate activation biases (*SI Appendix,* SMethod3). The estimated current densities of each subject were then averaged across all time points that were attributed to a given microstate in each condition. To evaluate whether the sources of the different microstates differed between the three conditions, t-tests across subjects were calculated for each solution point (*SI Appendix,* Table S1). Finally, the sources of a given microstate were averaged across subjects for each condition (*SI Appendix,* SFigs. 3-5).

## Supplementary Methods 1: Experimental Paradigm

Each participant underwent three sessions: i. an interview session, ii. an EEG recording session, and iii. an fMRI session. During the interview, all participants performed a classical autobiographical memory questionnaire (ABMQ) (1) in order to select vividly remembered images, which were then used for the EEG and fMRI sessions that included three distinct conditions (*Supplementary Figure,* Fig. S1): eyes-closed rest (6min), mentally retrieving personal past episodes (10min), and mental arithmetic operations (10min). The memory and math conditions comprised twenty 30-s trials each. On each trial, a personal image (e.g., photo of a participant with a birthday cake) or a calculation (e.g., 447-7=) was presented for 5sec, followed by a 20sec period of closed-eyes during which the participants retrieved the past event or continued to serially subtract/add the given number. After this period, there was a question: *“How much did you relive the original event?”* for the memory task and *“How much attention did you pay to the calculation?”* for the mental arithmetic task. The time allowed to read and respond to the question was 5sec. Participants answered by a button press (1=not at all to 4=fully). The ratings confirmed active involvement of our participants with high confidence ratings that were comparable between the EEG (M=3.2, SEM=0.12) and the fMRI (M=3.1, SEM=0.11) memory session as well as the EEG (M=3.5, SEM=0.12) and the fMRI (M=3.41, SEM=0.11) math session.

## Supplementary Methods 2: fMRI Acquisition and Analysis

### fMRI Acquisition

The fMRI sessions were conducted on an actively shielded Magnetom 7T head scanner (Siemens, Erlangen, Germany), equipped with AC84 head gradients (80mT/m max. gradient strength, 333 T/m/s max. slew-rate) and an 8-channel transmit/receive head loop array (Rapid Biomedical, Rimpar, Germany). During each paradigm, whole-brain functional data were acquired using a simultaneous multi-slice (SMS) 2D GE-EPI sequence, with TR/TE = 1000/22ms, α = 54°, 110×110 matrix size with 2.0 mm isotropic spatial resolution, 75 sagittal slices with 3× SMS acceleration and 1/3 field-of-view (FOV) CAIPI shift (1), 2× in-plane GRAPPA acceleration (anterior-posterior direction) and 7/8 partial Fourier undersampling. This protocol was designed to simultaneously provide fairly high temporal resolution (1 volume/s) and spatial resolution (2-mm voxel width), with whole-brain coverage. For each subject, an additional 5-volume scan was also performed with reversed phase encoding direction (posterior-anterior), for subsequent correction of susceptibility-induced EPI distortions. To aid spatial co-registration, T_1_-weighted anatomical data were acquired with a 3D MP2RAGE sequence (2) with TR/TI_1_/TI_2_/TE = 5500/750/2350/1.87 ms and 1 mm isotropic spatial resolution.

### fMRI Pre-processing

The fMRI data were processed using the FMRIB Software Library (FSL v5.0, Oxford, UK) combined with Matlab routines developed in-house (Mathworks, Natick MA, USA). Data pre-processing included motion correction (MCFLIRT tool, 6 degrees of freedom) (3), slice-timing adjustment (set to the middle of each TR, via linear interpolation), brain segmentation (BET tool) (4), Gaussian spatial smoothing (FWHM = 4mm) and temporal de-trending (100sec cut off). To reduce contributions from head motion, a set of 24 confound regressors derived from the motion parameter time courses were regressed out of the fMRI data by general linear model analysis. Each dataset was co-registered to the standard MNI space as follows: first, for each subject, the fMRI data were B_0_-unwarped using FSL-TOPUP (5) and brought to the subject’s anatomical space using FLIRT with boundary-based registration (6) (12 degrees of freedom). The co-registration to MNI space was then determined using the anatomical images, again through FLIRT (12 degrees of freedom).

### Fractional Amplitude of Low Frequency Fluctuations (fALFF) Analysis

Voxel-wise fALFF values were estimated for the pre-processed fMRI data of each subject and paradigm, in native space, as proposed by Zou et al. (7): the time series of each voxel was transformed to the frequency domain and the sum of amplitudes in the 0.01–0.08 Hz interval was divided by the sum of amplitudes of the full frequency band. The individual fALFF maps thus obtained for each paradigm and subject were then brought to MNI space for group analysis. The fALFF maps were compared between the memory and math conditions using paired T-tests across subjects, and statistical significance was determined using topological false-discovery rate (FDR) inference to correct for multiple comparisons (8) (cluster-forming threshold T > 2.20, FDR α = 5%).

### Functional Connectivity Analysis

The brain regions (clusters) displaying significant fALFF changes between the memory and math conditions were further considered for connectivity analysis. These regions were warped to each individual run’s native space, and the voxels belonging to each region were identified. The functional connectivity between each pair of regions was estimated as the average Pearson correlation value between the time course of every voxel in the first region with that of every voxel in the second region. The estimations were organized in a connectivity matrix, and also included an estimation for each region paired with itself, serving as a measure of functional homogeneity within the region.

## Supplementary Methods 3: EEG Acquisition and Analysis

### Pre-processing

EEG data were band-pass filtered between 1 Hz and 40 Hz using a noncausal filter (2nd order butterworth Low and High pass, - 12 db/octave roll-off, computed linearly forward and backward, eliminating the phase shift, and with poles calculated each time to the desired cut-off frequency). Infomax-based Independent Component Analysis (ICA) was applied to remove oculomotor and cardiac artefacts based on the channels with maximal amplitude, the topography, and time course of the ICA component.

### K-means clustering

A modified k-means clustering (1) was applied to the data of each subject and condition. Only maps at local maxima of the Global Field Power (GFP) entered the cluster analysis as they represent time points of highest signal-to-noise ratio (2). Cluster-analysis was applied in two steps: First on the data of each individual subject and condition, and then on the cluster maps derived from each subject within a condition. In order to determine the optimal number of clusters (both within and across subjects) 7 criteria have been used to evaluate independently the quality of each clustering. They were then merged together in order to derive a single synthetic meta-criterion. This improves the confidence in the right estimation of the optimal number of clusters, as compared to previous work relying on single criterion only (i.e. Cross-Validation criterion (3) or the Krzanowski-Lai Index (4)).

The following 7 criteria were taken from (3, 5–7):

*Gamma:* An adaptation of Goodman and Kruskal, based on concordant *vs.* discordant clustered pairs.

*Silhouettes*: Evaluation of the consistency of each cluster through its goodness of fit.

*Davies and Bouldin:* A function of the sum of the ratio of within-cluster to between-cluster separation.

*Point-Biserial*: A point-biserial correlation calculated between the distance matrix and a binary cluster index.

*Dunn*: An evaluation of the goodness of separation of all clusters.

*Krzanowski-Lai Index:* A ratio of the relative difference of the within clusters dispersion.

*Cross-Validation*: A modified version of the predictive residual variance.

The meta-criterion was defined as the median of all optimal number of clusters across all criteria (*Supplementary Figures,* Fig. S6). The meta-criterion calculation is implemented in the free academic software Cartool (https://sites.google.com/site/cartoolcommunity/). Note that this is an improved method of the earlier version that was used in (8).

### Back-fitting

The six group cluster maps of a given condition were fitted back to the original EEG of each subject, including all data points (not only GFP peaks) except for periods that were marked as artefacts. Back-fitting means that the spatial correlation between the six cluster maps and each individual data point was calculated and the data point was labelled with the cluster map that showed the highest correlation (winner takes all, see e.g. (9, 10)). Note that the polarity of the maps was ignored in this back-fitting procedure. Data points where none of the six maps reached a correlation higher than 50% were labelled as “non-assigned”. Once the whole recording was labelled, a temporal smoothing was applied by ignoring segments where a given cluster map was present for less than 4 time points (32ms) and the time points were split and assigned to the preceding and following cluster map. Two temporal parameters were then quantified for each recording of each subject: the mean duration a given cluster map was present without interruption, and the number of times per second a given microstate appeared, independent of the duration – termed “occurrence”(11).

### EEG source localization normalization

After the inverse solution matrix has been applied to the ongoing EEG at each time point, a standardization of the estimated current density across time was applied to each solution point. As a matter of fact, substantial variability of power is observed across solution points when localizing the sources on ongoing (non-averaged) EEG in individual subjects. These variations are supposed to come from geometrical and mathematical approximations that are done during the inverse matrix calculation. It is thus necessary to find a way to correct for this power variability, in order to correctly estimate the fluctuations of brain activity over time in individual subjects and to compare them between subjects (*Supplementary Figure,* Fig. S7).

The main idea here is to use the background activity of the norm of the inverse solution over time to estimate a baseline and a scaling factor for each solution point. In order to have a robust estimation, a large enough time sample should be used, preferably the whole recording as it was done in this study. Still, the correction factors can be satisfactorily computed on as little as a thousand time points, as long as no solution point remains in the same stable state more than half of the sampled time.

Here is a step-by-step description of this specialized standardization:

Given a 3D dipole (*sp*_*x*_, *sp*_*y*_, *sp*_*z*_) at a given solution point *sp*, we define *sp*_*χ*_ as the squared value of its norm (Equ. 1). It therefore follows a Chi-square distribution of degree 3 for the noisy part of the data. The Chi-square variable *sp*_*χ*_can be transformed into a normally distributed variable *sp*_*N*_ (Equ. 2) (12). Having now a normal distribution, it can be standardized into *sp*_*Z*_ by using the regular z-transform (Equ. 3). However, the values of μ and σ used for the z-transform have to be calculated only on the noisy part of the data - the background activity from the Chi-square i.e. the lowest values of the distribution. Hence μ is estimated from the left-most Mode of the *sp*_*N*_ distribution (Equ. 4). For the same reason, σ is estimated from the Median of Absolute Deviation (MAD), centred on the previously estimated μ, and computed only with the values below μ (Equ. 5). Implementation-wise, these estimators are computed multiple times on random sub-samplings of the data, and the two respective medians of all these estimators are finally taken.

Finally, because we started with positive data (the norm of a dipole), we also wish to end up with positive data, as to avoid any confusion due to having signed results. We define *sp*_*Z*+_ as *sp*_*Z*_ shifted by 3 standard deviations to the right, then divided by 3 so that the background mode is finally aligned to 1 (Equ. 6).

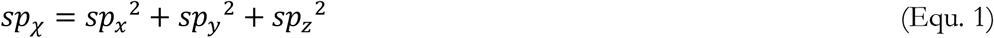

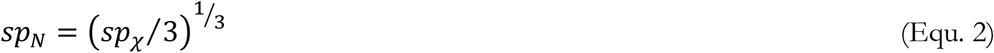

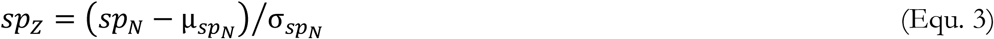

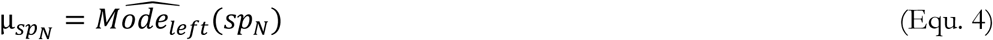

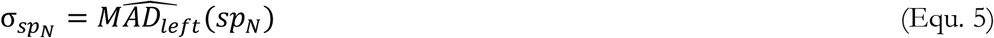

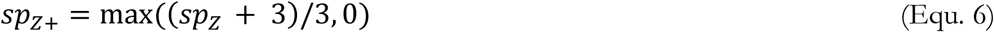

The result of this standardization procedure is comparable power of the current density across all solution points, and a normal distribution of the solution points (*Supplementary Figure,* Fig. S7).

## Supplementary Figures

**SFig. 1.**
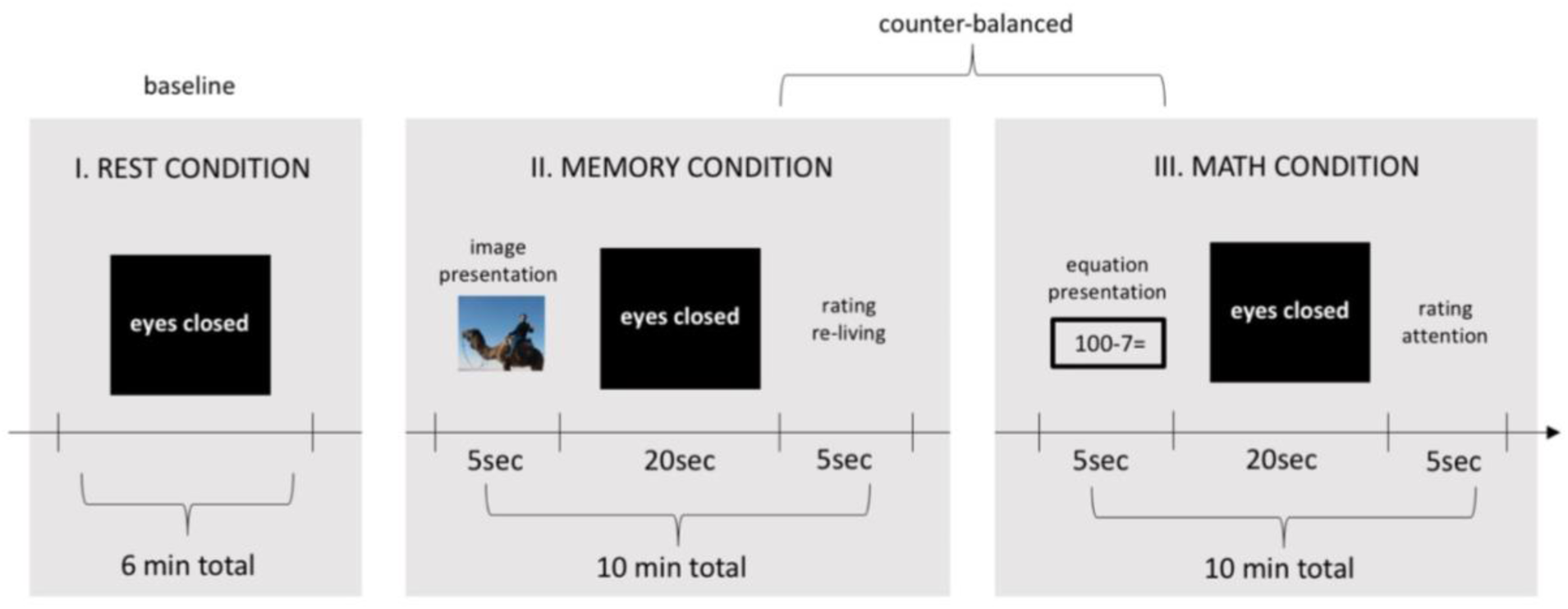
Experimental paradigm. 15 healthy participants (30.5±5.5years) took part in both high-density 64-channel EEG and 7T-fMRI sessions in three distinct conditions during eyes closed: rest (6min), mentally retrieving personal past episodes (10min), and mental arithmetic (10min). Before the scanning, all participants performed a classical autobiographical memory questionnaire (ABMQ)(28) in order to select vividly remembered images for the memory retrieval condition. Before each of the memory retrieval and mental arithmetic trials, participants saw a personal image/calculation for 5sec, after which they closed their eyes and had to either retrieve the event or continually subtract the numbers (20sec). After each trial, there were 5sec for a control question to answer: *“How much did you relive the original event?”* for memory condition and *“How much attention did you pay to the calculation?”* for the math condition (scale 1=not at all to 4=fully).

**SFig. 2.**
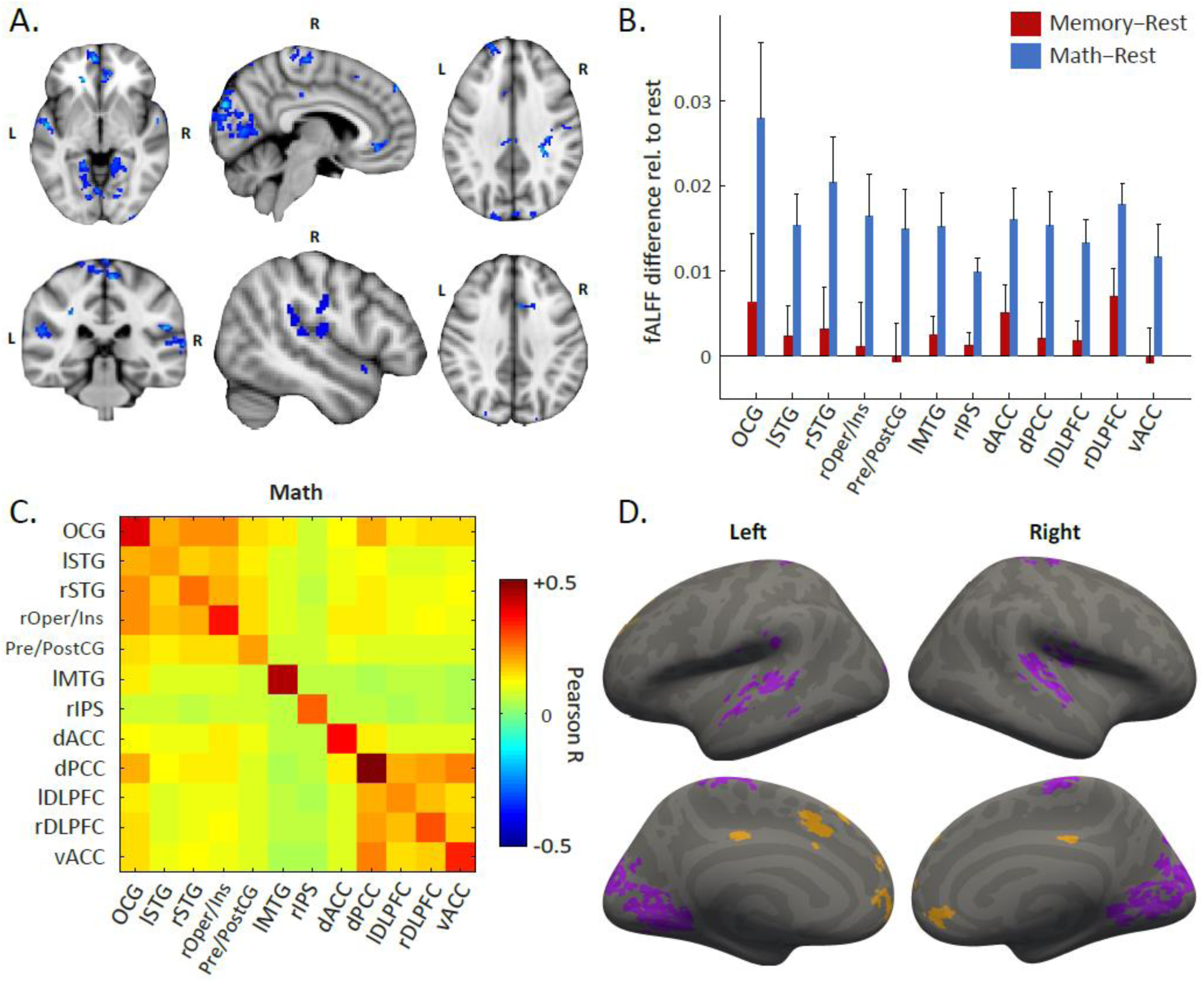
Math-specific changes in fMRI activity and fMRI functional connectivity networks. **A. fMRI fALFF analysis.** T-test of fALFF for math > memory condition revealed stronger activity in a set of brain areas. In the frontal lobe, significantly higher activity was found in the ventral and dorsal anterior cingulate cortex (vACC, BA24; dACC, BA32), and bilaterally in the dorsolateral prefrontal cortex (lrDLPFC, BA46). Areas in the temporal lobe included bilaterally the superior temporal gyrus (lrSTG, BA22), the left middle temporal gyrus (lMTG, BA21), and the right opercular cortex/insula (rOper/Ins, BA45). In the parietal cortex, significantly higher activity was found in the right intraparietal sulcus (rIPS, BA7) and the pre/post central gyrus (Pre/PostCG, BA3,1) and dorsal posterior cingulate cortex (dPCC, BA30). The occipital findings concerned the occipital gyrus bilaterally (OCG, BA19). T-score maths > memory threshold: T>2.20; FDR-corrected for multiple comparisons (5%). **B. Areas of stronger activity in the math, compared to memory condition.** In contrast to the above observation of areas de-activated during math (compared to the rest condition), none of the areas with significantly higher activity in math compared to memory were actually de-activated during the memory condition. **C. Connectivity analysis.** The connectivity analysis revealed two distinct networks within the regions that had displayed stronger activity during the math condition: network III included the OCG, lrSTG, rOper/Ins and Pre/PostCG, while network IV comprised the dACC, dPCC, lrDLPFC and vACC. Beyond their strong intrinsic connections, the two networks were also moderately connected to each other, unlike what was observed for the memory-related networks. The lMTG and rIPS regions displayed no relevant correlations with any of the two networks, or with each other. **D. Sub-networks of the connectivity analysis.** The two identified networks were also projected onto a surface template for better visualization.

**SFig. 3.**
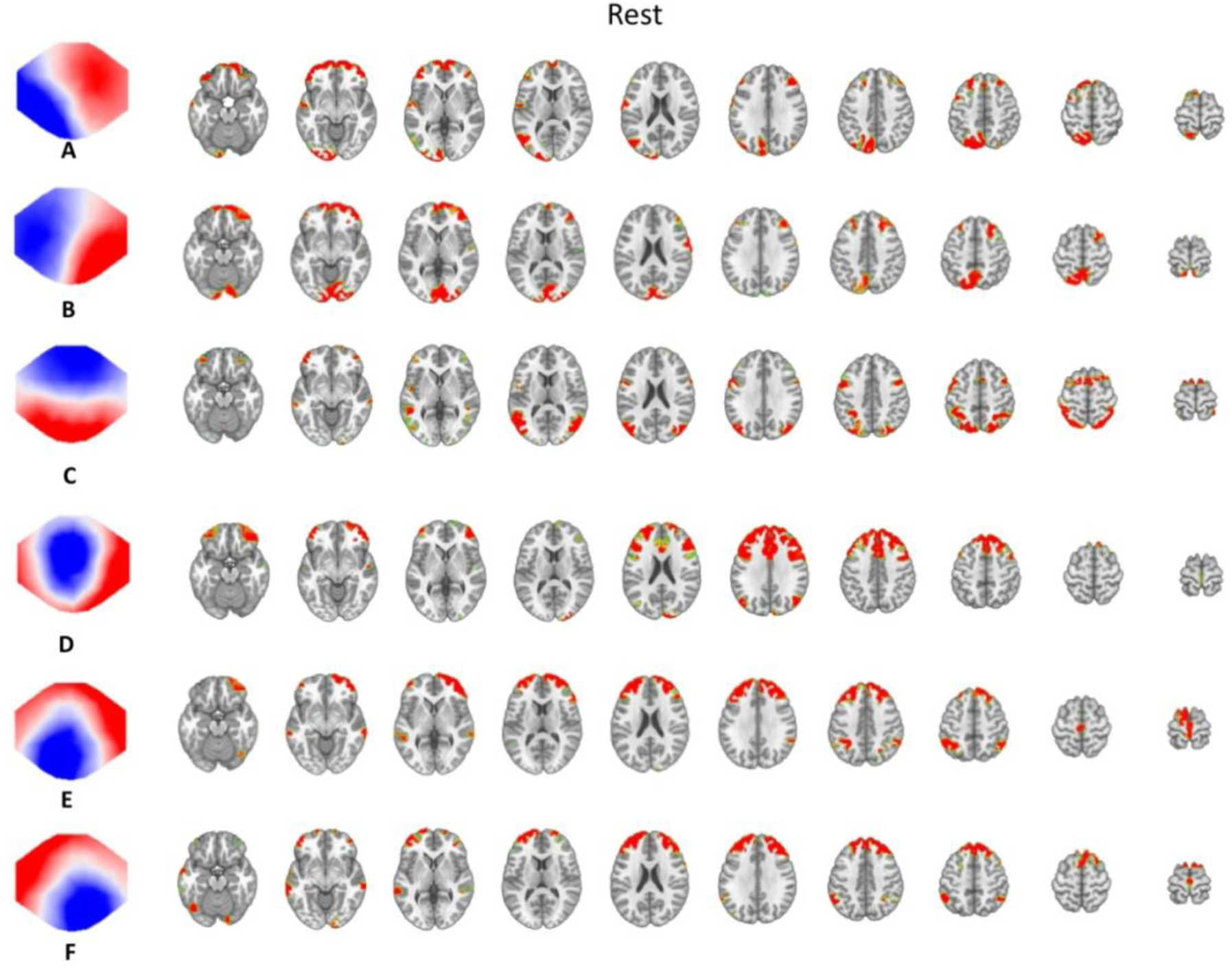
Source localization of the six microstates during the rest condition. Sources are averaged across all time points that were labelled with the corresponding map and averaged across subjects. Areas with activity above 95 percentiles are shown.

**SFig. 4.**
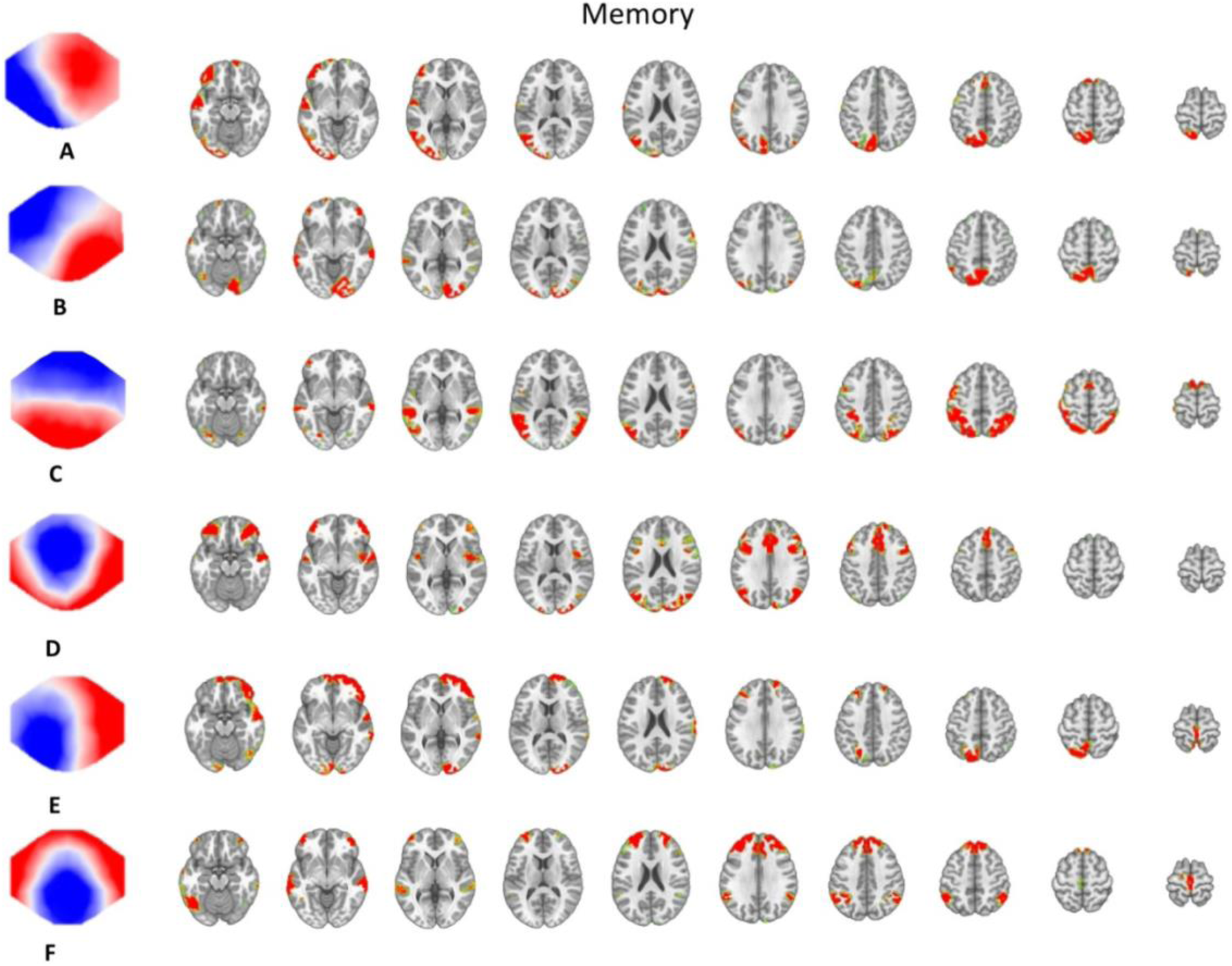
Source localization of the six microstates during the memory condition. Sources are averaged across all time points that were labelled with the corresponding map and averaged across subjects. Areas with activity above 95 percentiles are shown.

**SFig. 5.**
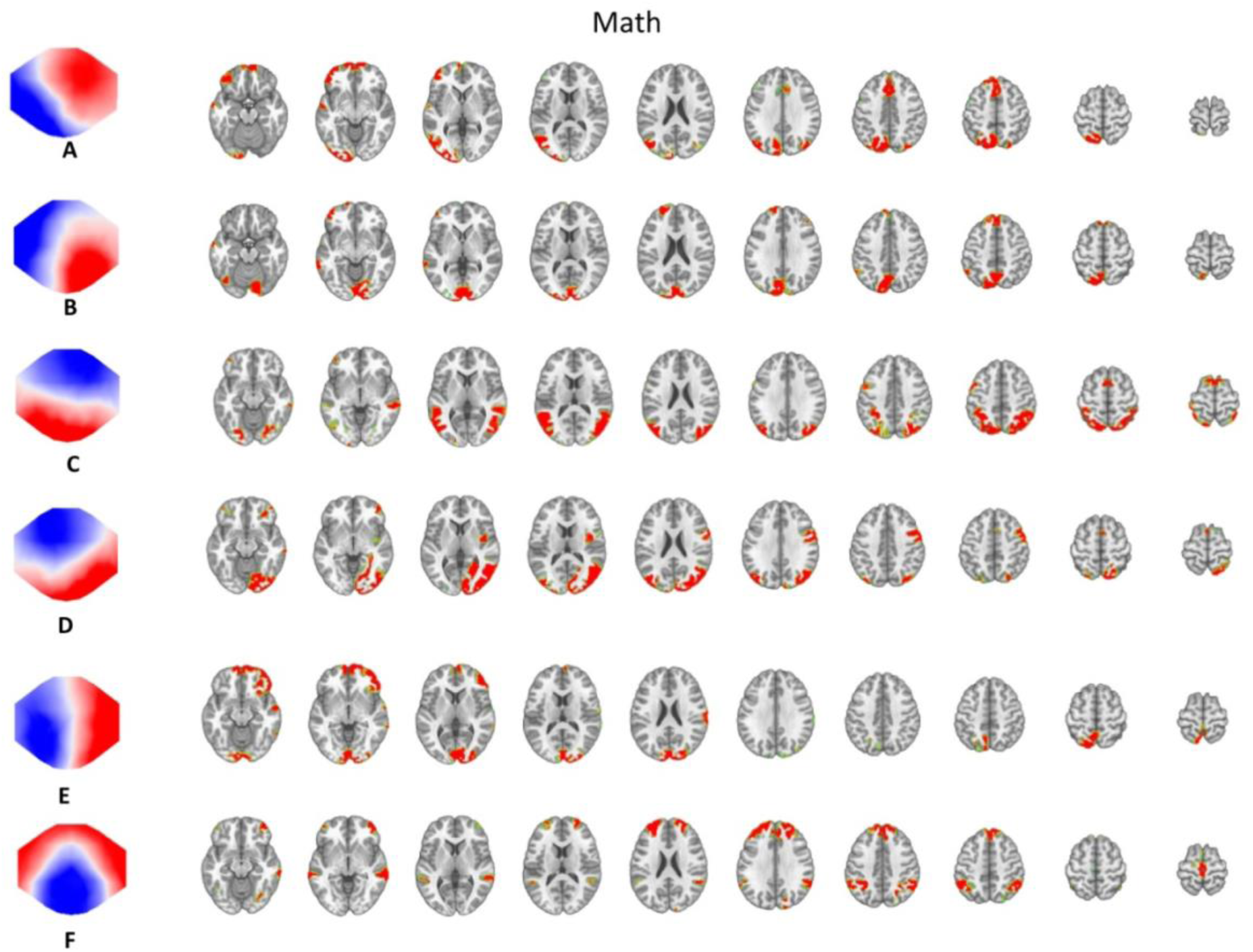
Source localization of the six microstates during the math condition. Sources are averaged across all time points that were labelled with the corresponding map and averaged across subjects. Areas with activity above 95 percentiles are shown.

**SFig. 6.**
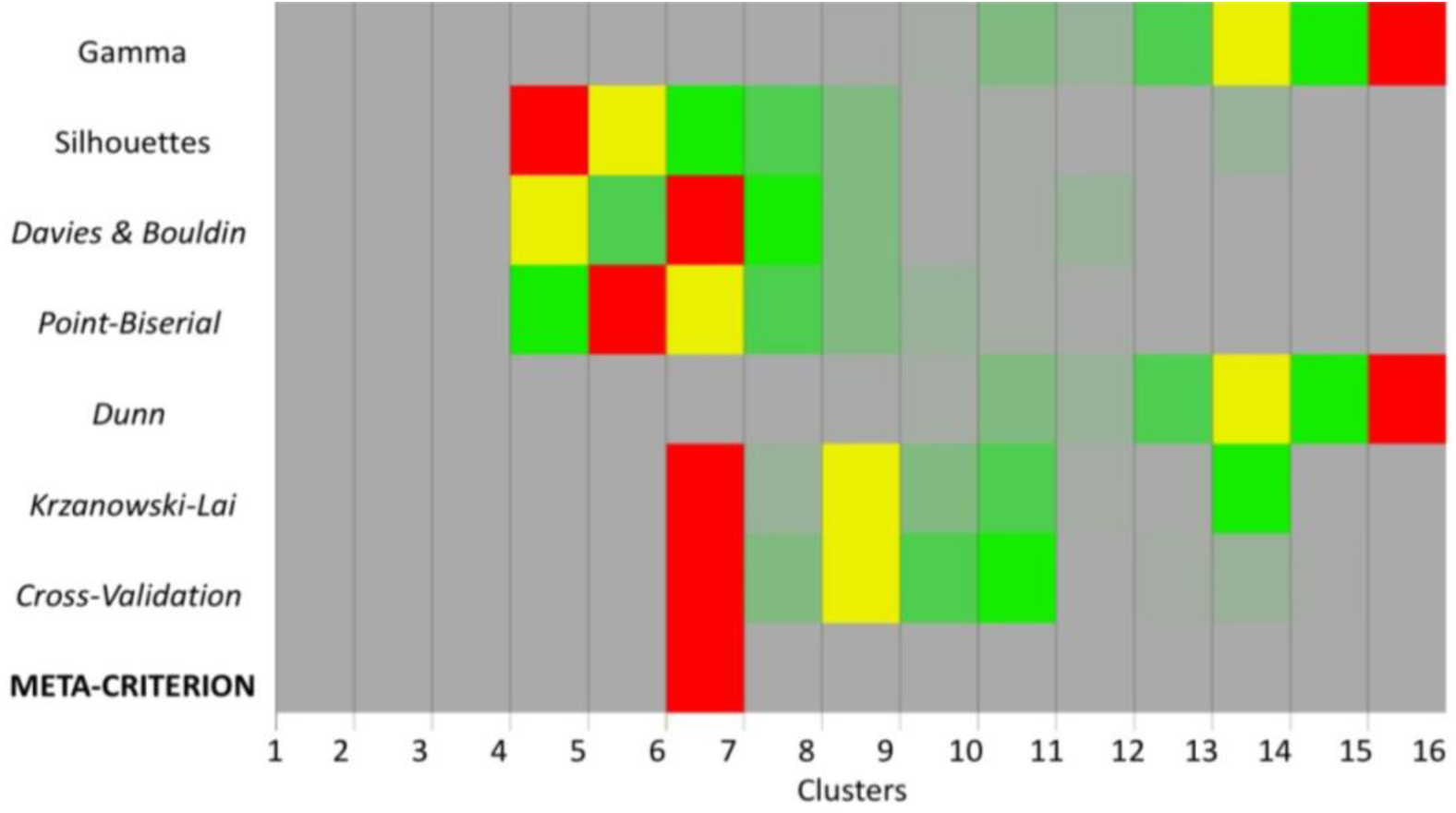
Criteria for evaluating the optimal number of microstate maps. 7 criteria (vertical axis) have been used to evaluate independently the quality of each clustering (horizontal axis). A red square points where a given criterion peaks, *i.e.* the optimal number of cluster for that criterion. The meta-criterion (last row) is defined as the median of all the optimal number of clusters across all criteria.

**SFig. 7.**
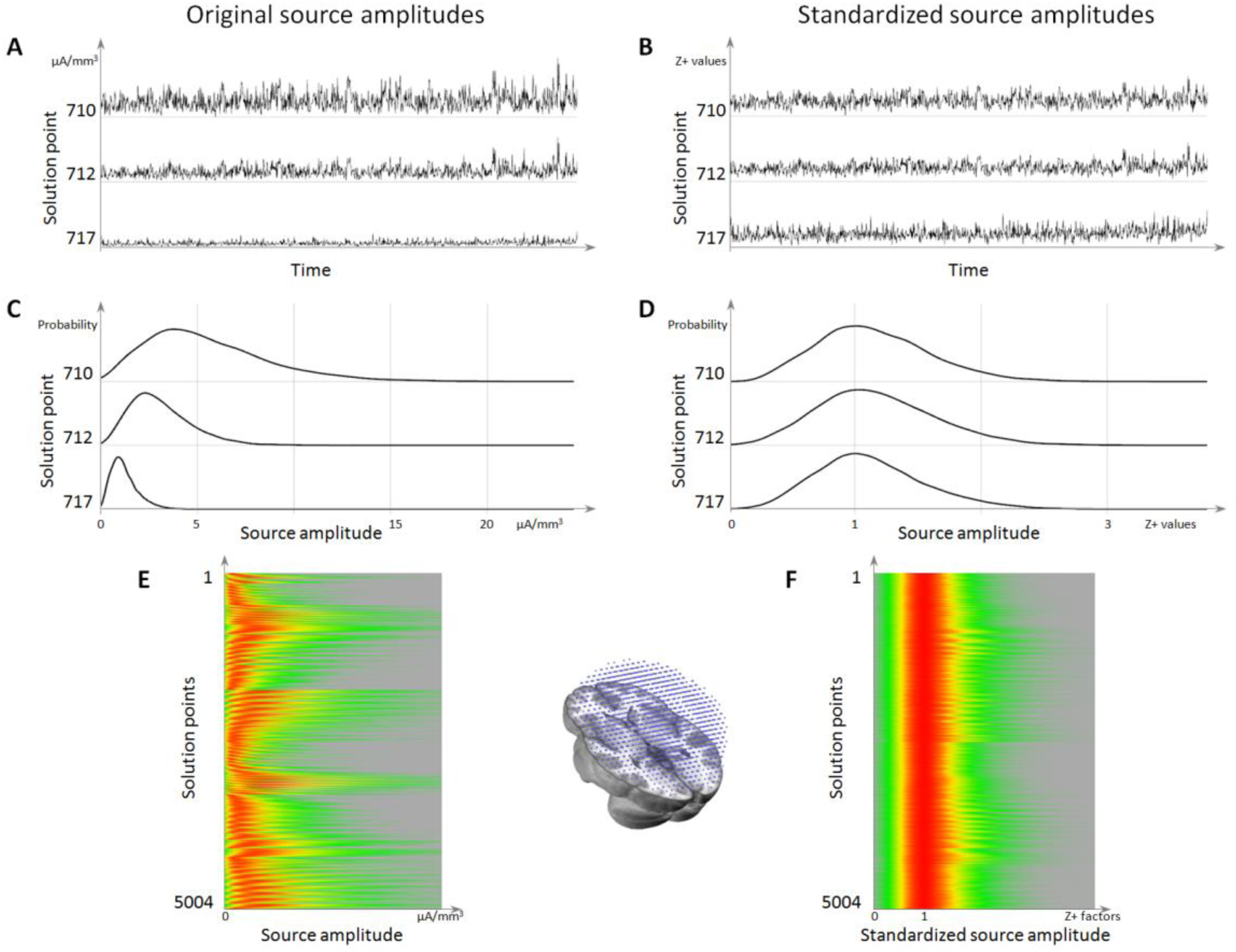
Source localization standardization. **A.** is the time series (horizontal axis) of 3 solution points (vertical axis), showing the difference in power range between them. **C.** is the histograms of these 3 solution points, showing that the background activity is the left-most mode of the distribution. **E.** is the histograms for all solution points (vertical axis), with the red color coding for the maximum probability. **B.** is the same time series as in **A** but after standardization, showing that the 3 solution points now have the same range. **D.** is the histograms after standardization, showing that all the background activity has been centered to 1. **F.** is all histograms after standardization, showing that all solution points now have a background range from 0 to 1, while retaining their respective highest activities.

**S Table1.**
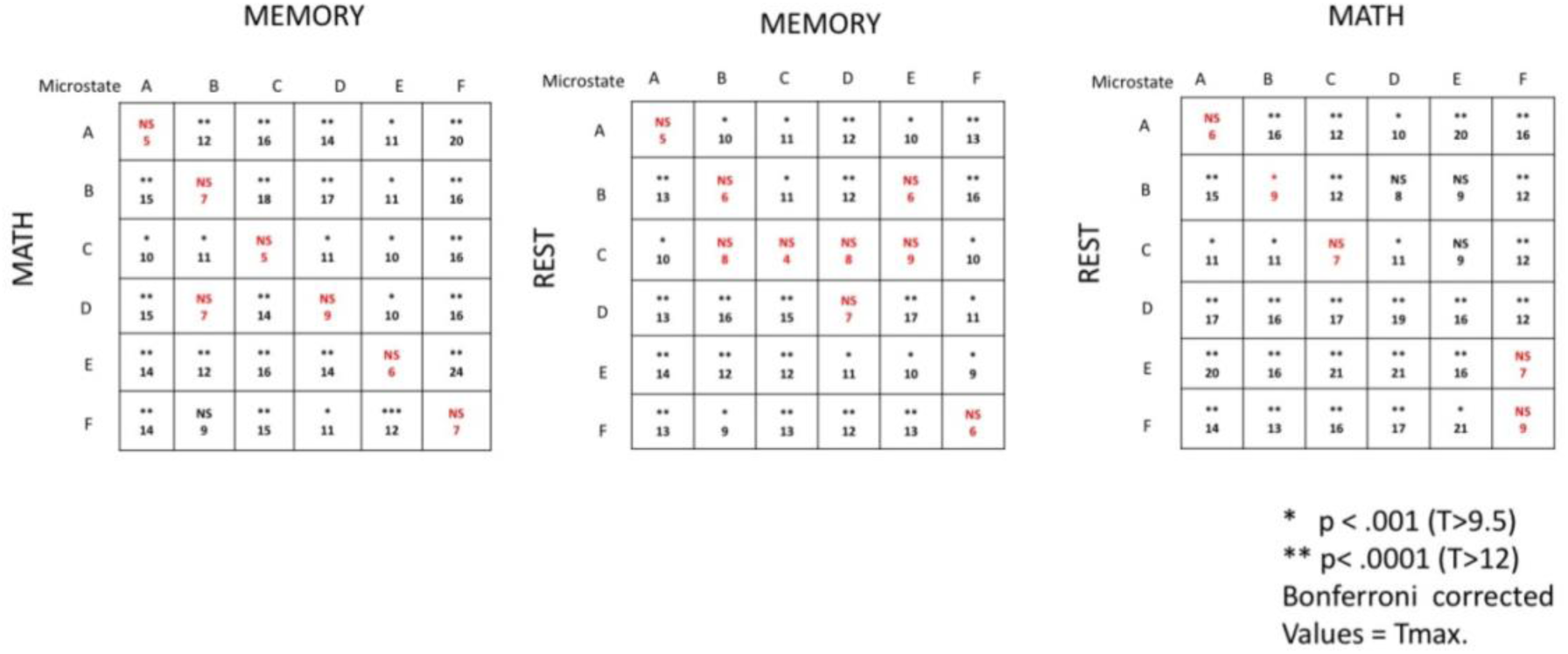
Result of the T-test across subjects of the sources derived for each microstate in each of the three conditions. Significance values and maximal T-values are given. Note that in general the sources of the microstates with the same label are not significantly different, while the sources of different microstates are. An exception is microstate D and E in the comparison of the math vs. the rest condition. In this case all significantly different solution points for microstate D were located in the superior frontal gyrus where stronger activity was found in the rest condition.

|  |  | MEMORY |  |  |  |  |  |
| --- | --- | --- | --- | --- | --- | --- | --- |
| MATH | Microstate | A | B | C | D | E | F |
|  | A | NS<br>5 | **<br>12 | **<br>16 | **<br>14 | *<br>11 | **<br>20 |
|  | B | **<br>15 | NS<br>7 | **<br>18 | **<br>17 | *<br>11 | **<br>16 |
|  | C | *<br>10 | *<br>11 | NS<br>5 | *<br>11 | *<br>10 | **<br>16 |
|  | D | **<br>15 | NS<br>7 | **<br>14 | NS<br>9 | *<br>10 | **<br>16 |
|  | E | **<br>14 | **<br>12 | **<br>16 | **<br>14 | NS<br>6 | **<br>24 |
|  | F | **<br>14 | NS<br>9 | **<br>15 | *<br>11 | ***<br>12 | NS<br>7 |
| REST | Microstate | A | B | C | D | E | F |
|  | A | NS<br>5 | *<br>10 | *<br>11 | **<br>12 | *<br>10 | **<br>13 |
|  | B | **<br>13 | NS<br>6 | *<br>11 | **<br>12 | NS<br>6 | **<br>16 |
|  | C | *<br>10 | NS<br>8 | NS<br>4 | NS<br>8 | NS<br>9 | *<br>10 |
|  | D | **<br>13 | **<br>16 | **<br>15 | NS<br>7 | **<br>17 | *<br>11 |
|  | E | **<br>14 | **<br>12 | **<br>12 | *<br>11 | *<br>10 | *<br>9 |
|  | F | **<br>13 | *<br>9 | **<br>13 | **<br>12 | **<br>13 | NS<br>6 |

|  |  | MATH |  |  |  |  |  |
| --- | --- | --- | --- | --- | --- | --- | --- |
| REST | Microstate | A | B | C | D | E | F |
|  | A | NS<br>6 | **<br>16 | **<br>12 | *<br>10 | **<br>20 | **<br>16 |
|  | B | **<br>15 | *<br>9 | **<br>12 | NS<br>8 | NS<br>9 | **<br>12 |
|  | C | *<br>11 | *<br>11 | NS<br>7 | *<br>11 | NS<br>9 | **<br>12 |
|  | D | **<br>17 | **<br>16 | **<br>17 | **<br>19 | **<br>16 | **<br>12 |
|  | E | **<br>20 | **<br>16 | **<br>21 | **<br>21 | **<br>16 | NS<br>7 |
|  | F | **<br>14 | **<br>13 | **<br>16 | **<br>17 | *<br>21 | NS<br>9 |

\* p < .001 (T>9.5)
\*\* p < .0001 (T>12)
Bonferroni corrected Values = Tmax.

